# The *Doublesex* sex determination pathway regulates reproductive division of labor in honey bees

**DOI:** 10.1101/314492

**Authors:** Mariana Velasque, Lijun Qiu, Alexander S. Mikheyev

**Author notes:** Corresponding author: Alexander S Mikheyev.

## Abstract

Eusociality, the ultimate level of social organization, requires reproductive division of labor, and a sophisticated system of communication to maintain societal homeostasis. Reproductive division of labor is maintained by physiological differences between reproductive and sterile castes, typically dictated by pheromonal queen fertility signals that suppress worker reproduction. Intriguingly, reproduction and pheromonal signalling share regulatory machinery across insects.The gene *Doublesex* (*Dsx*) controls somatic sex determination and differentiation, including the development of ovaries and secondary sexual characteristics, such as pheromonal signalling. We hypothesized that this regulatory network was co-opted during eusocial evolution to regulate reproductive division of labor. Taking advantage of the breakdown in reproductive division of labor that occurs in honey bees when workers commence to lay eggs in the absence of a queen, we knocked down *Dsx* to observe effects on ovary development and fertility signal production. As expected, treated workers had lower levels of egg yolk protein, for which *Dsx* is a cis-regulatory enhancer in other insects, and greatly reduced ovary development. Also as expected, while control workers increased their levels of pheromonal fertility signals, treated workers did not, confirming the role of *Dsx* in regulating pheromone biosynthesis. We further found that *Dsx* is part of a large network enriched for regulatory proteins, which is also involved during early larval development, and upregulated in queen-destined larvae. Thus, the ancient developmental framework controlling sex specification and reproduction in solitary insects has been exapted for eusociality, forming the basis for reproductive division of labor and pheromonal signalling pathways.

**Significance statement:** Complex social insect societies rely on division of reproductive labor among their members. Reproductive individuals (‘queens’) suppress ‘worker’ reproduction using pheromonal fertility signalling. We show that an ancient regulatory network that controls specification of sex and secondary sexual characteristics in solitary insects, has been co-opted for both both pheromonal signalling and ovary inactivation in honey bees. In addition, this network is also active during caste specification that takes place during the first few days of larval life. These results show that pheromonal signalling and ovary development share a common regulatory framework, potentially explaining why fertility signalling is ‘honest.’ Furthermore, they show that higher levels of biological complexity can arise by rewiring and elaborating ancestral gene regulatory networks.

## Introduction

Eusociality has evolved multiple times across animal lineages, reaching its pinnacle in the social insects. It features ‘queen’ and ‘worker’ castes, which perform reproductive and provisioning tasks, respectively. These castes differ physiologically, with workers forming the disposable ‘somata’ of these societal superorganisms, and queens the ‘germlines.’ The organization of complex social insect societies depends on sophisticated communication systems that modulate reproductive output and other aspects of colony function. Because chemical communication generally evolves rapidly (1, 2), and the evolutionary origins of castes are much older, most studies have addressed these aspects of social function separately. However, recent work has demonstrated that queen pheromones are structurally conserved across lineages that independently evolved eusociality, and that pheromonal communication may be pleiotropically linked with caste physiology (3–6). These observations suggest that the two may be controlled by a common regulatory network dating back to solitary ancestors.

Ancestrally, solitary females performed both reproductive and provisioning behaviors, which became decoupled into the queen and worker castes of eusocial colonies (the “ovarian ground plan hypothesis”) (7). This transition may have involved elaboration of a common ancient “genetic toolkit” across different eusocial lineages(8). Building on this hypothesis, over a decade of studies have identified a large number of candidate genes, which include members of nutrient-sensing and juvenile hormone signalling pathways, storage proteins, and a DNA methyltransferase (reviewed in ref. 9). However, the pleiotropic interaction between gene regulatory networks, which control caste and pheromonal signalling, has not received much attention until relatively recently, though hormone levels and fertility signalling were shown to be coupled in ants and wasps (4, 5).

Queen pheromones are honest signals of an individual’s reproductive status (reviewed in ref.10). On one hand, such honesty could lead to the optimal function of a colony, as it allows workers to replace under-performing queens. On the other hand, honest signalling may enable policing of worker reproduction by nestmates. For example, in honey bees under a range of circumstances, workers may activate ovaries in an attempt to reproduce independently by laying male-destined eggs (11). However, the chemical signatures of laying workers become more queen-like (12), and they risk execution by their nestmates (13, 14). So why should workers advertise their fertility? One possibility is that, unlike many other traits of the solitary ancestor that became decoupled in queens and workers, pheromones and fertility remain functionally linked due to gene regulatory constraints(6).

This link is most plausibly seen in the honey bee, where sex and queen pheromones have been studied most extensively. The queen mandibular pheromone (QMP), acts as both a queen pheromone, suppressing worker reproduction, and as a sex pheromone, attracting males during nuptial flights (15). Surprisingly, both QMP biological functions may be deeply conserved, as QMP can inhibit ovary activation and attract males in fruit flies (16, 17). While the molecular mechanisms by which QMP acts as a queen pheromone remain poorly understood, these findings suggest an ancestral function involving both sexual attraction and fertility signalling by the same suite of chemicals.

A functional link between fertility and sexual signalling should generally be adaptive in solitary insect females, which have little incentive to hide their reproductive status from potential mates. Indeed, across insect orders, *doublesex* (*Dsx*), the transcription factor that regulates sexual differentiation in somatic tissues and also controls female sex pheromone production and reception (18, 19), further binds to *cis*-regulatory enhancers of egg yolk proteins (20–22). Based on these observations we hypothesized that *Dsx* and associated genes may have been co-opted during evolution of reproductive division of labor. This model could parsimoniously explain both reproductive division of labor though worker ovary inactivation, and the concomitant evolution of pheromonal communication systems among a variety of social insects.

We experimentally tested this hypothesis in honey bees. In the absence of a queen, reproductive division of labor breaks down as young workers activate ovaries and commence to lay male-destined eggs, while producing a queen-like blend of mandibular pheromone components (12). We predicted that RNAi-mediated *Dsx* knockdown would inhibit worker ovary development, specifically by reducing vitellogenin (Vg) expression, because a well-established cis-regulatory link between *Dsx* and egg yolk protein (specifically Vg) exists (20–22). We also predicted that *Dsx* knockdown should reduce the amount of queen-like mandibular pheromone produced by workers. We find support for both hypotheses, illustrating the central role of *Dsx* in the reproductive division of labor. We then used this experimental perturbation to characterize its role in the gene regulatory network of honey bees. Intriguingly, genes involved in this network also play a role during caste determination in young larvae. These data suggest that honey bee eusociality exapted ancient core developmental networks, specifically those involved in sex specification.

## Results

Supplementary figures and tables along with the R code necessary to generate them are hosted on http://mikheyevlab.github.io/dsx-rnai/.

### Validation of *Dsx* knockdown and *Vg* response

We sequenced twenty libraries split evenly among treatment and control groups on an Illumina HiSeq 2500 sequencer at the OIST Sequencing Center (SQC). The experiment yielded 3.5×10^7^ ± 4.7×10^6^ (s.d.) RSEM-mapped single-end reads per library. The overall fit of observed *vs.* expected spike-in controls transcripts explained 87% of the variance, indicating adequate technical performance. The fit did not increase at subsequent abundance cutoffs; therefore, we used data for downstream analyses without additional filtration. We detected significantly lower *Dsx* expression in RNAi-treated worker abdomens using both the RSEM/edgeR and Kallisto/Sleuth pipelines (Figure 1A). Levels of *Vg* were also significantly lower in bees where *Dsx* was knocked down, and were strongly correlated with those of *Dsx*, as predicted (Figure 1B).

**Figure 1.**
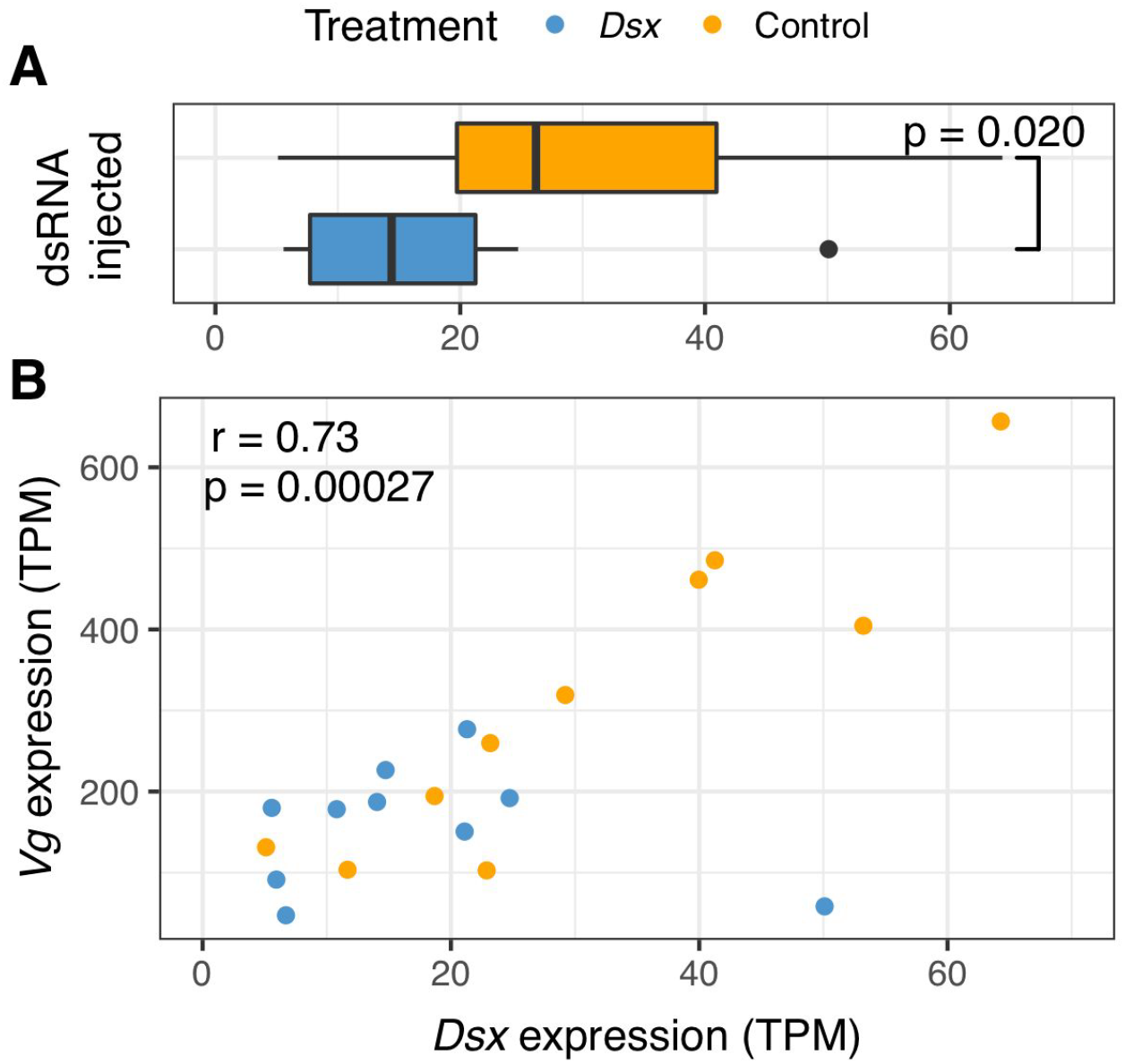
*Dsx* knockdown confirms the regulatory link between *Dsx* and *Vg* in honey bees. (A) RNA-seq confirmed that mean expression levels of *Dsx* were reduced 43.3% relative to GFP-injected controls (measured in Transcripts Per Million (TPM) mapped reads). (B) As predicted, expression of *Dsx* and *Vg* were tightly linked overall, and *Vg* expression levels were significantly lower in *Dsx-*knockdown bees (one-tailed p = 0.0082). *Vg* is an egg yolk precursor, and its levels in workers are strongly socially repressed in queenright colonies (69). These data support the central role of *Dsx* in honey bee physiology, in part by direct control of *Vg*, a key gene involved in coordinating diverse aspects of social organization (49, 50).

### Ovary and pheromonal activation

Queens and workers can produce overlapping sets of mandibular components, but major axes of variation successfully separate the two castes, and some components serve to discriminate fertility levels (reviewed in ref. 23). The first two principal component axes summarizing pheromonal variation in our data, explained 74% of the variance (49.5% and 24.9%, respectively). The first principal component (PC1) was correlated with 10-HDAA, 10-HDA, 9-HDA and HOB (r = 0.27, 0.25, 0.20 and 0.25, respectively) (Table S1). 10-HDA and 10-HDAA are worker-typical components (24, 25). Therefore, PC1 is most correlated with axes of worker mandibular gland pheromone variation. The second principal component (PC2) was strongly correlated with MGP components 9-HDA, 9-ODA and HVA (r = 0.26, 0.37 and 0.17, respectively). Greater levels of 9-HDA and HVA are associated with fecundity in queens (26), and 9-ODA is the ‘canonical’ queen mandibular pheromone in the bouquet used to attract males (15). Therefore higher values along the PC2 axis indicate more queen-like mandibular gland pheromone profiles.

Pheromonal profile and ovary activation are often linked (26, 27), but the specific relationship between pheromone profile composition and ovary activation stage is unknown. Therefore we simultaneously analyzed both phenotypic responses via a cumulative link principal component regression model and ovary activation stage as the response variable. To account for correlations among mandibular pheromone compounds (Figure S2), first two principal components were used as predictors, along with *Dsx* knockdown treatment, and interactions (Figure 2; Table S2). *Dsx* knockdown resulted in lower ovary development when compared to the non-target control gene (Z = 3.1, one-tailed p = 0.0011) and more worker-like scores on the PC2 axis (Z=-2.2, one-tailed p=0.015). Furthermore, there was a significant interaction between these two explanatory variables (Z = 2.3, p = 0.021), which meant that *Dsx*-knockdown disrupted the relationship between ovary activation level and the amount of pheromone produced. This can be seen in Figure 2, where control workers show the typical positive association between ovary activation and queen-like mandibular pheromone production, and *Dsx*-knockdown workers show the opposite pattern.

**Figure 2.**
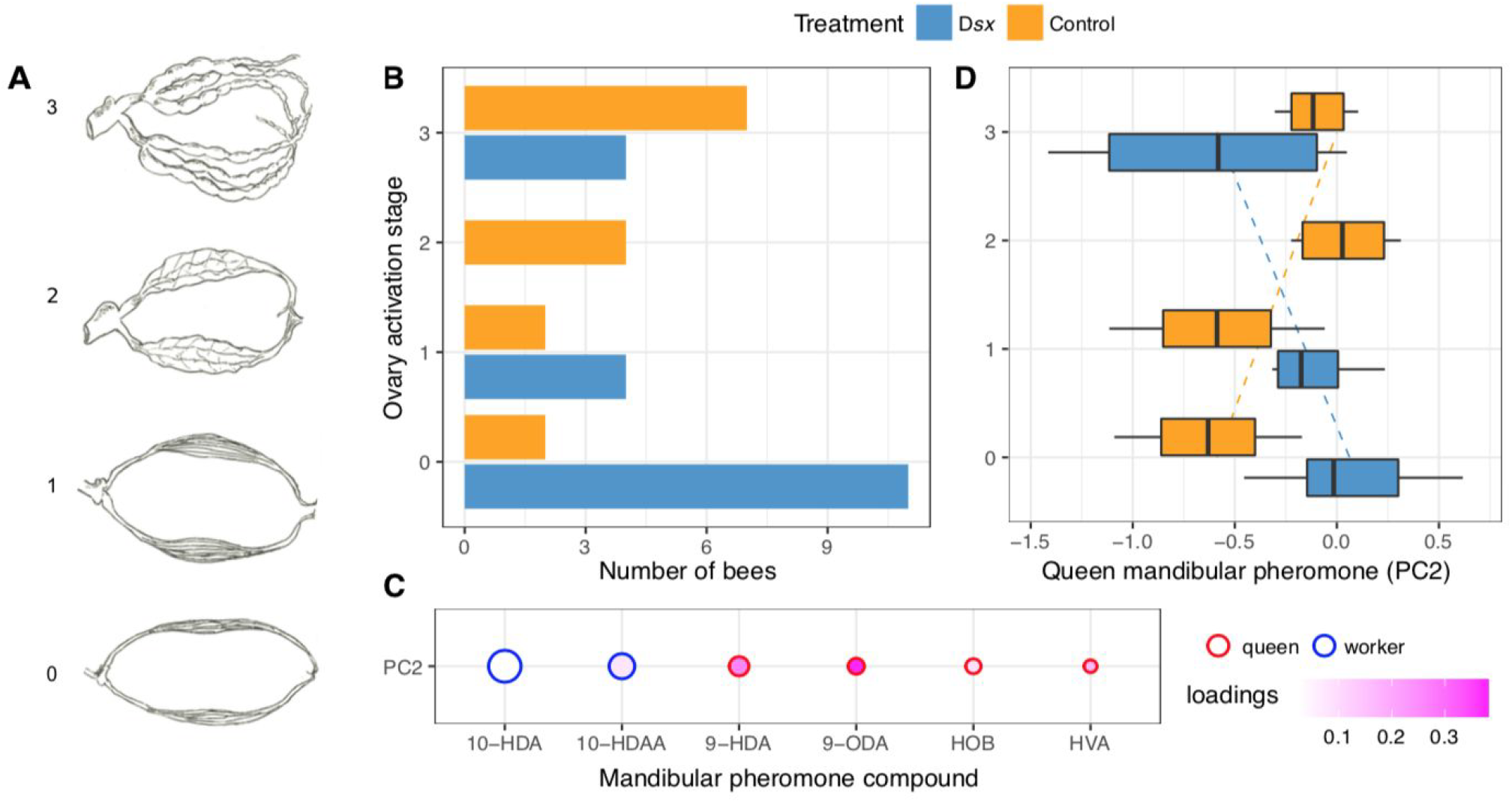
*Dsx* knockdown decreases ovary development and production of queen-like mandibular gland components. (A) Worker ovary activation stage. Stage 0 represents underdeveloped ovaries typical of young workers. Stage 1 represents enlarged ovaries. Stage 2 represents ovaries that have begun to develop, containing small oocytes, and Stage 3 denotes developed ovaries with eggs. (B) Ovary activation was significantly lower in *Dsx*-injected samples *vs.* controls, indicating that *Dsx* controls ovary development, as predicted (Z = 3.1,one-tailed p = 0.0011). (C) Higher values along the second principal component axis corresponded to more queen-like mandibular gland profiles (red circle outlines) (26, 70). Circle size represents the relative total amount of each component. (D) *Dsx* knockdown affected the relationship between ovary activation stage and levels of queen-like mandibular pheromones. Box plots show medians and ranges capturing 75% and 95% of the variation, with dashed trend lines showing robust linear model fits to help visualize trends. Our major prediction, that levels of queen mandibular pheromones (QMP) would be lower in treated bees was confirmed (Z = −2.16, one-tailed p = 0.0011). Typically, when workers start to lay eggs, they produce more mandibular pheromone components (12). However, the relationship between ovary activation stage and pheromonal production, and whether *Dsx* alters this relationship are unknown. Therefore we included an interaction term (Treatment×PC2) in the model to investigate it. The slope of the relationship was indeed different for *Dsx*-treated workers, also suggesting that *Dsx* regulates QMP production, but perhaps not as a simple on-off switch (Z = 2.3, p = 0.021). Given that *Dsx* controls both fertility signalling and ovary development in other insect orders *(18, 20–22)*, our data show that *Dsx* retained these ancestral functions, although the significant interaction term suggests the potential for additional regulatory elaboration over pheromone production.

These changes were not likely due to differences in vitality between the two treatment groups, as they did not differ significantly in mortality (Z = 0.083, p = 0.93, Table S3). Overall mortality levels were similar to other RNAi studies (e.g. (28, 29) and within the range expected for the diet (30).

### Identifying the *Dsx*-responsive gene regulatory network

Weighted gene co-expression analysis (WGCNA) takes advantage of correlations between gene expression patterns of genes across libraries to identify ‘modules’ of genes showing similar expression profiles (31). WGCNA has been shown to reconstruct protein-protein interaction networks with reasonable accuracy, based solely on gene expression data (32, 33). Therefore, WGCNA allowed us to examine the gene regulatory network surrounding *Dsx.* In particular we took advantage of the concept of ‘module membership’, which is the overall connectedness of a gene to other members of the same network. More important genes tend to have greater membership.

WGCNA identified a network containing *Dsx* and 966 other genes, out of a total of 13,811 (Table S4). Genes in this network were generally over-expressed in control bees, and their module membership was strongly correlated with the log2 fold-count of control *vs.* treated gene expression, *i.e.*, genes most involved in the network showed the greatest responses to *Dsx* knockdown (Figure 3). This module also contained *Vg*, as would be expected, due to its tight regulation by *Dsx*. It also contained another well-known gene that was experimentally shown to affect caste, the DNA methyltransferase *Dnmt3 (34)* (Table S5). Within the co-expression module, *Dsx* was part of a tightly connected network of transcription factors and other regulatory proteins (Figure S3). Overall, the module was enriched for gene ontology terms associated with regulation, particularly of transcription, and signalling (Table S6). Genes in this module were upregulated in control bees, and genes that were more tightly connected to other genes in the module were more likely to respond to *Dsx* knockdown (Figure 3A). Therefore, sensitivity to *Dsx* knockdown is a core property of this module.

**Figure 3.**
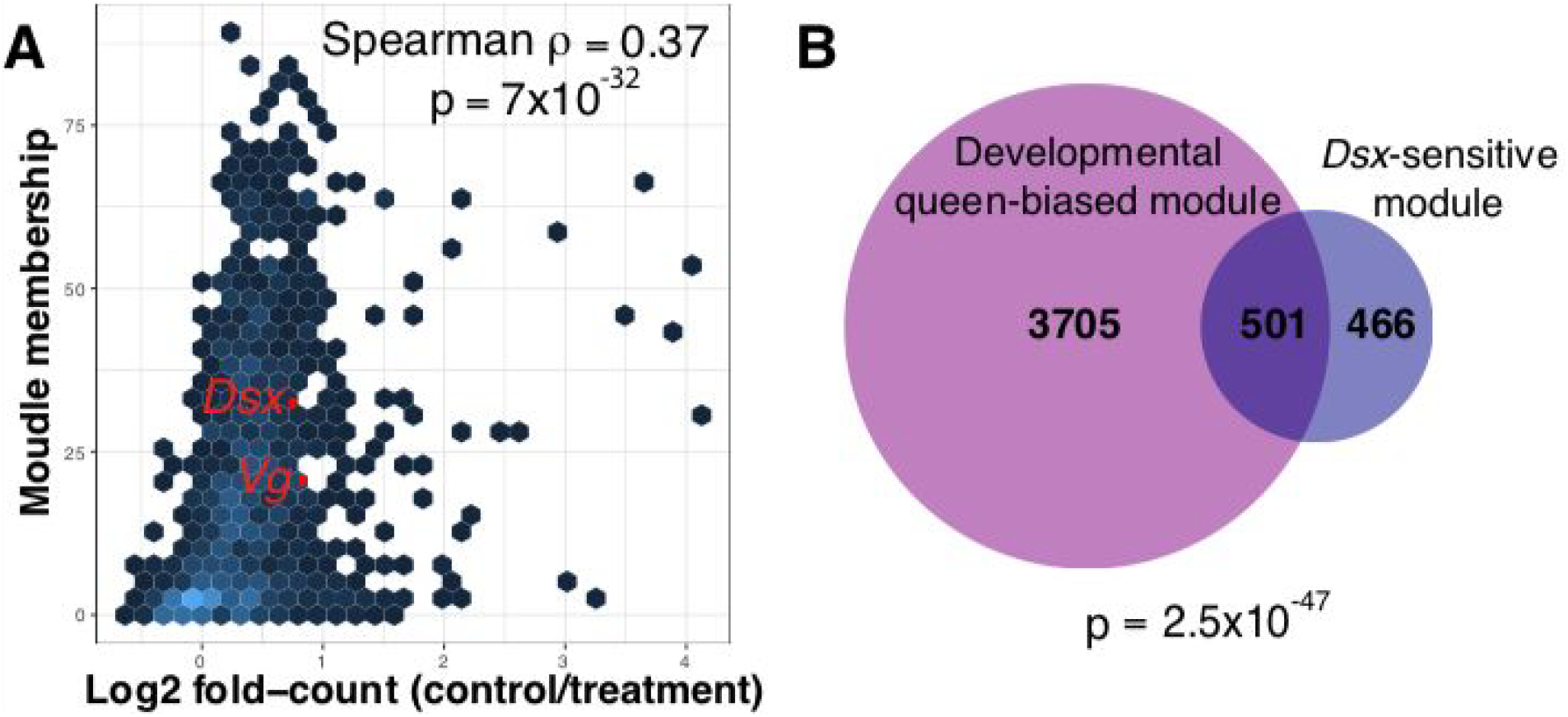
Co-expression networks surrounding *Dsx* are associated with reproductive division of labor in adults and during larval caste differentiation. (A) The *Dsx-*responsive module identified from experimental data showed a strong correlation between module membership and differential expression log-fold count (control *vs.* knockdown). Point color indicates the number of data points in each hexagon (up to a maximum of 20 in the lightest-colored ones). Most fold-count values are positive, indicating that these were generally upregulated in control bees (*i.e.*, in reproductively active workers). Module membership measures the extent to which a gene’s expression is correlated with that of other genes in the module. Genes most deeply integrated into the module were also most affected by the experimental treatment, indicating that responsiveness to *Dsx* knockdown is a core feature of this module. As expected, in addition to *Dsx*, this module also contained *Vg,* because expression of both genes is correlated (Figure 1B), and *dnmt3*, which has been shown to affect caste determination (34). Figure S1 shows an interactive version of panel A. (B) The *Dsx*-responsive responsive module that was experimentally identified in adults significantly overlapped a large *Dsx*-containing module comprising queen-biased genes that is active during early larval development. This suggests that regulatory mechanisms involved in reproductive differentiation in adults and larvae significantly overlap and may share a common core.

#### Does *Dsx* play a role during larval caste differentiation?

In honey bees caste differentiation takes place during early larval development. We asked whether mechanisms identified for maintaining reproductive division of labor in adults also play a role at that stage. While a previous study looking at gene expression during larval caste specification found no differential expression of *Dsx (35)*, in our experimental perturbation, we found that the gene-level signal was relatively weak (Figure 1A), possibly due to tissue-specific expression of *Dsx (36)*, but the network-level signal and its effects were dramatic (Figure 3A). Therefore we asked whether the *Dsx*-sensitive module present in adults might also play a similar role in the context of larval caste determination.

We examined module preservation in two ways. First, we used an integrated statistic (Z_summary_) in the WGCNA package (31), which tests for preservation of a given module in expression data. According to this test, the *Dsx-*responsive module detected in our experiment was strongly preserved in whole-body larval gene expression data (35), meaning that it was also co-expressed in 2 and 4 day old queen- and worker-destined larvae (p = 9.9×10^−7^). In 4-day old larvae, which have committed to their caste-specific developmental trajectories, these genes were generally upregulated in future queens (rho = 0.083, p = 0.015). Second, we performed WGCNA on the same data set, and identified a large module that contained *Dsx* (Table S7). Membership in this module was likewise correlated with queen development (rho = 0.049, p = 0.0023). Half of the genes (50.5%) from the experimentally identified *Dsx*-responsive adult module, were found in this developmental module (Figure 3B). These results suggest that similar gene regulatory networks play a role in both adult and larval caste differentiation, and are involved in inducing reproductive phenotypes in both cases.

#### *Major royal jelly protein* also responds to *Dsx* knockdown

A number of genes have been experimentally shown to affect larval caste determination in honey bees (reviewed in ref. 9). From that list, only *Vg* and *Dnmt3* were part of the co-expression module containing *Dsx*. Though part of a different module, *major royal jelly protein* (*Mrjp1*) was also upregulated in control bees (log_2_ fold-count = 2.78, p = 0.0036). This gene shows caste-specific expression and its monomeric form is a key dietary component inducing queen development (37).

## Discussion

While primary mechanisms of insect sex determination are diverse, they always converge on the *Dsx* pathway, which involves the regulation of various sexually dimorphic traits (38). Insects have co-opted *Dsx* for a variety of different functions, ranging from sexual ornamentation in beetles (39–41), to butterfly wing patterning (42). Importantly, *Dsx* regulates ovary development and the production of sex pheromones (18, 20–22). Consequently, the *Dsx* pathway provides an ancient gene regulatory framework coupling two building blocks of eusociality – differential fertility and pheromonal signalling. We thus hypothesized that it has also been co-opted during the evolution of eusociality. Experimental knockdown of *Dsx* confirmed that it regulates ovary activation, likely by regulating the egg yolk precursor *Vg* (Figure 1), and in pheromonal signalling in adult workers (Figure 2). We identified a *Dsx*-responsive gene co-expression module that is enriched in gene ontology terms associated with biological regulation (Figure 3A). This module is also preserved in larval gene expression data, and is specifically associated with the queen-destined developmental trajectory (Figure 3B). These data suggest that in honey bees, reproductive division of labor evolved by taking advantage of existing sex-specific developmental regulatory networks.

Social evolution may have taken advantage of a ‘genetic toolkit’ consisting of ancient gene regulatory networks that have been re-wired for social living (8, 43). Although many studies have provided support for this hypothesis, the components comprising this toolkit and how they interact remain unclear (9, 44, 45). Putative genes in the genetic toolkit that have been identified to date are enriched in members of nutrition and juvenile hormone-signalling pathways (reviewed in ref. 9), all of which act downstream of *Dsx* during development. For example, *Vg* is a key toolkit genes, and under its direct *Dsx* control. Other genes that have been experimentally shown to affect caste, such as *Dnmt3* and *Mrjp1* also responded to *Dsx* knockdown suggesting that they interact with it either directly or indirectly. Studies on horned beetles have shown that *Dsx* interacts with nutritional levels to produce alternative phenotypes, *i.e.*, the size of horns (40, 46). This likely also happens in honey bees, since *Vg* is sensitive to nutritional state in a variety of insects, including honey bees (47, 48). Furthermore, *Dsx* can indirectly control other aspects of colony function via *Vg*, which has many coordinating effects on social organization (49, 50).

*Dsx* is a nexus in insect development and physiology, interacting with a wide range of other regulatory proteins (38) (Table S6). Highly connected genes are expected to experience evolutionary constraints as a result of pleiotropic interactions (51, 52). Thus, interconnectedness of *Dsx* may impose constraints on how traits under its control evolve, since they are coupled to a conserved ancestral gene regulatory network. This functional constraint could explain why fertility signalling by queens is honest, and even why ‘cheating’ reproductive workers advertise their behavior despite the risk of execution (53, 54). The hypothesis that signalling and ovary activation are pleiotropically linked and therefore constrained to be honest, been proposed previously with experimental data from ants and wasps, though with an emphasis on interactions with juvenile hormone (4–6). Our data are consistent with these observations since *Dsx* directly regulates *Vg*, which is integrated into the juvenile hormone and insulin signalling pathways (55). Therefore, honest signalling may be a byproduct of the proximate mechanisms exapted for eusociality, rather than inherently adaptive in all cases.

If *Dsx* is central to reproductive division of labor in honey bees, why has it not been picked up by other studies as a core gene involved in caste determination, even those that specifically tested for its differential expression (e.g., (35))? Expression of *Dsx* is highly tissue-specific (36, 41, 56). Aggregating data at the whole body or body part level is common practice in social insect gene expression studies, but this likely increases the signal-to-noise ratio. This hypothesis is supported by the observation that the only study to date that has detected significant differences in *Dsx* in honey bees analyzed tissue-specific expression (36). Furthermore, because *Dsx* interacts with many other genes, its levels may be constrained within fairly narrow ranges, making differences harder to detect. For example, though we attempted to maximize *Dsx* variability in our experiment, it varied only by an order of magnitude (Figure 1B), though that was sufficient to induce phenotypic effects (Figure 2). Future work should examine fine-scale changes in the expression of *Dsx* across honey bee tissues and over time to identify developmental hotspots that may be associated with caste determination and other aspects of social function.

## Conclusion

Building on prior knowledge of the extensive role of *Dsx* in insect development and polyphenism, we experimentally showed that it participates in a core gene regulatory network that is involved in reproductive division of labor in honey bees and that can indirectly control many aspects of social function by regulating *Vg* levels (49). It may also play a role in epigenetic reprogramming, as it occurs in the same module as *dnmt3*, a DNA methyltransferase known to affect caste (34). We also found that this network is associated with caste differentiation during early larval development. Furthermore, a recent study found that *Dsx* is differentially expressed among castes in an ant (57), and coupling between ovary development and reproductive signalling has been reported in ants and in wasps (4, 5), suggesting that exaptation of the sex determination pathway may have been widespread for independent origins of eusociality. A deeper understanding of the regulatory network surrounding *Dsx* and comparative studies should provide novel insights into the mechanisms, evolutionary potential, and constraints of eusocial evolution.

## Acknowledgements

We would like to thank Alejandro Villar of the OIST mass spectrometry center for carrying out the pheromone derivatization and for help with the interpretation of mass spectra. We also thank Jarol Chen for her drawings. We express appreciation to Steven D. Aird, Luke Holman, Armin Moczek, and Michael Warner for comments on the manuscript. We are grateful to Luke Holman for discussion and suggestions over the course of the study. This work was funded by OIST subsidy funding and by JSPS KAKENHI grants 16H06209 and 16KK0175 to ASM.

## Materials and Methods

### Laboratory experiments

Experimental procedures were performed blind. dsRNA *Dsx* and control solutions were labeled with distinct colors during microinjection, and their identities were not revealed during acquisition of pheromonal profiles and ovary activation data to prevent subconscious bias.

#### dsRNA Synthesis

We designed a pair of primers for the female-specific F2 isoform of *Dsx* (NCBI ID NM_001134935.1), which differs from the other *Dsx* isoform that is expressed in both sexes (NCBI ID NM_001134936). Primers were fused with the T7-promoter sequence (underlined) at their 5’ end(DSX-forward: <u>TAATACGACTCACTATAGGG</u>TTCTTCGGTCCCTCAACCAC; DSX-reverse: <u>TAATACGACTCACTATAGGG</u>GTCTGTGGCAAATGGGTGAC). We used a similar method to design the Green Fluorescent Protein (GFP) primer (GFP-forward: 5’-<u>TAATACGACTCACTATAGGGCGA</u>AGTGGAGAGGGTGAAGGTGA; GFP-reverse: <u>TAATACGACTCACTATAGGGCGA</u>GGTAAAAGGACAGGGCCATC).

We extracted total RNA from a 14 day old queen using TRIzol^®^ and synthesized cDNA as described by Aird *et al*. (58). We used the cDNA as the template to amplify the target portion of the *Dsx* gene and the pET6Xhn-GFPuv vector (Clontech) to amplify the *GFP* gene. We purified PCR amplicons by solid phase reversible immobilization with 19% PEG. The amplicons were then used as templates for dsRNA synthesis using the MEGAscrip kit (Ambion). The synthesized dsRNA products were purified using MEGAclear kit (Ambion) and eluted with nuclease-free water. The quantity and quality of dsRNA were evaluated using NanoDrop 2000C (Thermo Scientific) and Agilent RNA-6000 Pico kit, respectively.

#### Microinjection

We collected brood combs from *Apis mellifera ligustica* colonies from the apiary of the Okinawa Institute of Science and Technology (OIST), Okinawa, Japan. We used brood frames from six different colonies collected on October 23th and November 6th 2017 and incubated overnight at 35 ° C and 70% humidity (59). The next day, we collected approximately 300 newly emerged workers and randomly mixed them (150 were used on October 24th and 150 on November 7th 2017). Prior to injection, we immobilized bees by cooling them at 4 ° C for 1-2 minutes and fixed on beeswax plates using two crossed pins. In this position, we inserted the needle between the 3rd and 4th tergite (at the side of the abdomen) and injected newly emerged honey bees with 1 μL dsRNA solution (N=31) of the target gene DSX or 1μL dsRNA solution (N=31) of a non-target control gene green fluorescent protein (GFP), a non-honey bee gene. We diluted all dsRNA used in this experiment to 1ng/μL, as our pilot study shown a higher mortality rate when bees were injected with higher concentrations. Different microneedles and Microloader pipette tips (Eppendorf, cat. no. 5242 956.003) were used for each bee.

After the injection, we kept workers on the wax plates until their recovery and discarded all bees showing signs of haemolymph leakage. We housed experimental bees individually in cages containing 20 nurse bees (from a different experimental colony) and 5 newly emerged bees (from the same cohort) for 10 days at 35°C and 70% humidity. Studies with honey bees indicate that experimental cages can mimic the effects of a queenless colony (60), stimulating ovary development. Moreover, as royal jelly has a high nutritive value, promoting higher rates of ovary development, we also fed honey bees with 50% royal jelly mixed in honey and water (30).

#### Assessment of worker ovary development

On the 11th day after microinjection, we dissected surviving experimental bees (30). Heads were removed, immediately frozen and stored at −80 ° C for further pheromonal analysis (see below). To quantify ovary activation, we scored dissected ovaries on a scale from 0 to 3, as described by (59), with 0 being used for underdeveloped ovaries (without distinguishable oocytes), 1 for ovaries containing visible oocytes, 2 for ovaries containing sausage-shaped oocytes, and 3 when they had a fully developed egg. To prevent RNA degradation, honey bees were dissected submerged in RNA Later^®^ (Ambion), then frozen and stored at −80 ° C.

#### Analysis of mandibular gland pheromones

On the 11th day after microinjection, honey bee heads were frozen immediately after decapitation. They were transferred to 1 mL borosilicate glass tubes, inserted into a 2-mL Eppendorf tubes containing 400 μL of chloroform and 5 μL of internal standard solution (composed of 1 mg of octanoic acid, 1 mg of tetradecane in 4 mL dichloromethane) and stored for at least 24 h at −30 ° C. Prior to gas chromatography, the sample was divided in half (to be stored as a backup for further analysis) and the other half (200 μL) was evaporated until near dryness with a gentle nitrogen stream. The residue was redissolved in 30 uL BSTFA, pyridine at 2:1 ratio and incubated at 70 ° with shaking for 1 h. One μL of that solution was injected into the GC/MS. MGP concentration was estimated as: 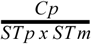, where Cp is the peak of the MGP compound, STp is the peak area of the internal standard, and STm is the internal standard concentration.

#### RNA-seq library preparation

From each treatment group, we selected 10 honey bees (20 total, 10 honey bees with lower and 10 bees with higher ovary activation scores). RNA was isolated using standard TRI-zol^®^ Reagent (Life Technologies) procedure, except that RNA was precipitated in the presence of 0.35 μL of glycogen (20 μg/μL concentration). Total RNA from each bee was diluted in 10 μL of nuclease free water. Quantity of RNA was examined with NanoDrop 2000c spectrophotometer (Thermo Scientific) and quality with Argument 2100 bioanalyzer (Agilent Technologies). cDNA synthesis, amplification, and preparation of RNA-Seq libraries was performed using the protocol described by Aird *et al*. (58), including the addition of ERCC92 spike-in controls. Libraries were sequenced on an Illumina HiSeq 2500, and raw reads were deposited into DDBJ under BioProject accession number PRJDB6980.

### Data analysis

All statistical analyses and results can be viewed on http://mikheyevlab.github.io/dsx-rnai/. We summarize them here in brief. We present two-tailed p-values throughout, except for the four statistical tests where we predicted specific directional effects in response to treatment in knockdown *vs.* control bees: levels of *Dsx* and *Vg*, and in levels of ovary activation and queen-like pheromone components (61).

#### RNA-seq analysis

The NCBI Annotation Release 103 of the honey bee (Amel_4.5) genome was used for differential gene expression analysis. Our goal was fourfold: (a) to validate *Dsx* knockdown, (b) to test for corresponding *Vg* decrease in knockdown bees, (c) identify the network of genes co-expressed with *Dsx*, and (d) examine the extent to which this network is involved in larval caste differentiation using data from He *et al.* (35) We employed two alternative approaches for differential gene expression analysis: a more traditional pipeline (RSEM) using read mapping and an alignment-free method (Kallisto) (62–64). We used edgeR for differential gene expression of RSEM data, and Sleuth for Kallisto data (65, 66). They gave the same results for *Dsx* and *Vg* levels, and we proceeded with the former pipeline for weighted gene co-expression analysis (WGCNA) to identify the *Dsx*-responsive module (31).

We conducted our analysis in adult bees because we were interested in the effects of *Dsx* on both ovary development and pheromonal signalling. However, we were also interested in determining whether similar mechanisms may be involved in larval caste determination. We tested for preservation of the *Dsx*-responsive module in the extensive data set from He *et al.* (35), who compared transcriptional differences between worker- and queen-destined larvae during the first four days of development.

#### Pheromonal level and ovary activation analysis

To the best of our knowledge, the relationship between ovary activation stage and mandibular gland profile has not been investigated. Therefore, we were interested in testing the relationship between *Dsx* knockdown, pheromone production, and ovary activation state in a joint analysis. Because ovary activation is an ordered factor variable, it was best suited as a response variable for a cumulative link model regression analysis. However, levels of mandibular gland components are not independent, as they share pathways (67), and can be highly correlated (Figure S2). To account for multicollinearity we conducted a principal component regression (68). The number of principal components was chosen by adding them stepwise in order of the amount of variance they explained, until the overall model fit ceased to improve, according to likelihood ratio tests. This resulted in two principal components as explanatory variables, which happened to explain a preponderance of the variance and also had biologically significant interpretations, being correlated with worker and queen axes of pheromonal variation. Furthemore, to ensure that effects were not due to concomitant differences in vitality, we also compared mortality between knockdown and control bees using a binomial mixed model.

